# Large field of view fluorescence imaging of microfluidic devices with a tandem-lens macroscope^†^

**DOI:** 10.1101/2025.10.11.680838

**Authors:** Daniel A. Mokhtari, Ali Lashkaripour, Polly M. Fordyce

## Abstract

Microfluidic devices enable high-throughput sample processing with remarkable parallelization and miniaturization. While fluorescence microscopy provides a convenient method for reading out signal from microfluidic assays, commercially-available microscopes impose a fundamental tradeoff between temporal resolution, spatial resolution, and numerical aperture (NA). Spatially tiled imaging enables high-resolution and high-NA imaging over a large area but reduces temporal resolution. Conversely, low magnification, low NA imaging captures large areas in one shot, but typically sacrifices spatial resolution and fluorescence sensitivity. To address this, we introduce an automated transfluorescence tandem-macro-lens optomechanical system (macroscope) capable of sensitive, multi-channel fluorescence imaging over a very large field of view (34 mm diameter, 740 mm^2^), with resolution determined by the sensor pixel size. We demonstrate bright-field resolution of low-micron features and detection of low-to mid-nanomolar concentrations of common fluorophores within microfluidic device channels. To demonstrate the utility of this macroscope, we image enzyme turnover within valved microfluidic devices (the HT-MEK system, for High-Throughput Microfluidic Enzyme Kinetics) and achieve >50-fold increased temporal resolution over common commercial instruments while maintaining high sensitivity. This macroscope imaging solution costs substantially less than commercially available alternatives, providing a powerful new imaging approach for microfluidic applications requiring sensitive and rapid wide-field fluorescence imaging.

---

Commercial epifluorescence microscopes are attractive turn-key solutions for imaging microfluidic devices because they offer ecosystems of easily upgradeable parts, robust interfaces and mount points, and built-in support for different objectives, filters, and stages. However, these systems can be simultaneously overbuilt and under-optimized for microfluidic applications. ^1^ For example, many commercial epifluorescence systems are engineered for stable nanometer-scale resolution (*e*.*g*. live cell imaging via confocal microscopy or super-resolution imaging), yet microfluidic applications involving beads, droplets, chambers, or wells often require imaging with modest (low micron) spatial resolution. Commercial systems are also typically limited to fields of view (FOVs) less than 100 mm^2^ (2X or higher magnification), while many microfluidic systems span regions of interest (ROIs) much larger, necessitating tiled imaging that limits temporal resolution. ^2,3^ While lower magnification objectives (*e*.*g*., 1X objectives) are commercially available, they tend to yield poor sensitivity due to low numerical aperture (NA), produce images with significant aberrations, and are limited in maximum FOV by the rest of the optical train. Thus, commercial fluorescence microscopes are simultaneously expensive and limiting for many microfluidic applications. ^4^

Examples of microfluidic applications that could benefit from high temporal resolution imaging include HT-MEK (HighThroughput Microfluidic Enzyme Kinetics) and *k*-STAMMP (kinetic Simultaneous Transcription Factor Affinity Measurements via Microfluidic Protein Arrays), which use valved microfluidic devices to express, purify, and quantify kinetic reaction rates for >1,500 protein variants in parallel. ^5–7^ These assays use fluorescence timecourse imaging to monitor kinetic processes within a large (18×18 mm) array of chambers containing features as small as 25 μm. When originally reported, these assays employed a Nikon Ti epifluorescence microscope to image the entire device via tiled acquisition of ~35–50 images at each time point, thus limiting temporal resolution to the time required to acquire all tiled images and return to the starting position (~30–75 s). This slow image acquisition limited the study of fast biophysical processes (*e*.*g*., rapid enzyme catalysis, binding kinetics, or diffusion and transport).

Tandem-lens epifluorescence macroscopy, a large FOV fluorescence imaging method originally developed for imaging mouse cortex with high temporal- and spatial-resolution, offers a potential solution. ^8^ In this system, two lenses are infinity-focused and assembled face-to-face to create an approximate “infinity space” between them that can accommodate optical elements such as filters and dichroic mirrors; the effective magnification depends on the ratio of the lenses’ focal lengths. Tandem-lens configurations have previously been leveraged to image large FOVs, including circular FOVs of ø9 mm^8^, as well as rectangular areas 12.6×10.5 mm^9^, 13.4×11.3 mm^10^, and 15×10 mm^11^. In addition to a large FOV, tandem-lens systems can provide very high NA (*e*.*g*., 0.36–0.58) while minimizing aberration. ^8–10^ However, FOVs of tandem macroscopes in epifluorescence configurations are constrained by clipping produced by the slightly divergent light exiting the objective lens and the large gap between the two lenses needed to accommodate a dichroic in the lens’ approximate “infinity space.” ^8,10^

Here, we adapt tandem-lens macroscopy to enable sensitive and rapid fluorescence imaging of large ROIs for microfluidic applications at significantly lower cost than common commercial fluorescence microscopes. Specifically, we present a tandem-lens macroscope that can: (1) capture a large FOV of hundreds of mm^2^, (2) automatically switch between filters for multi-channel fluorescence imaging, (3) capture images with high spatial resolution (minimum resolvable features of <10 μm) and fluorescence sensitivity at >1 frame per second (FPS), (4) illuminate a large FOV uniformly, (5) maintain a flat focal plane across the FOV, (6) position samples with high precision over a large travel distance, and (7) be motorized for automation (Fig. 1A and Table S1).

**Fig. 1:**
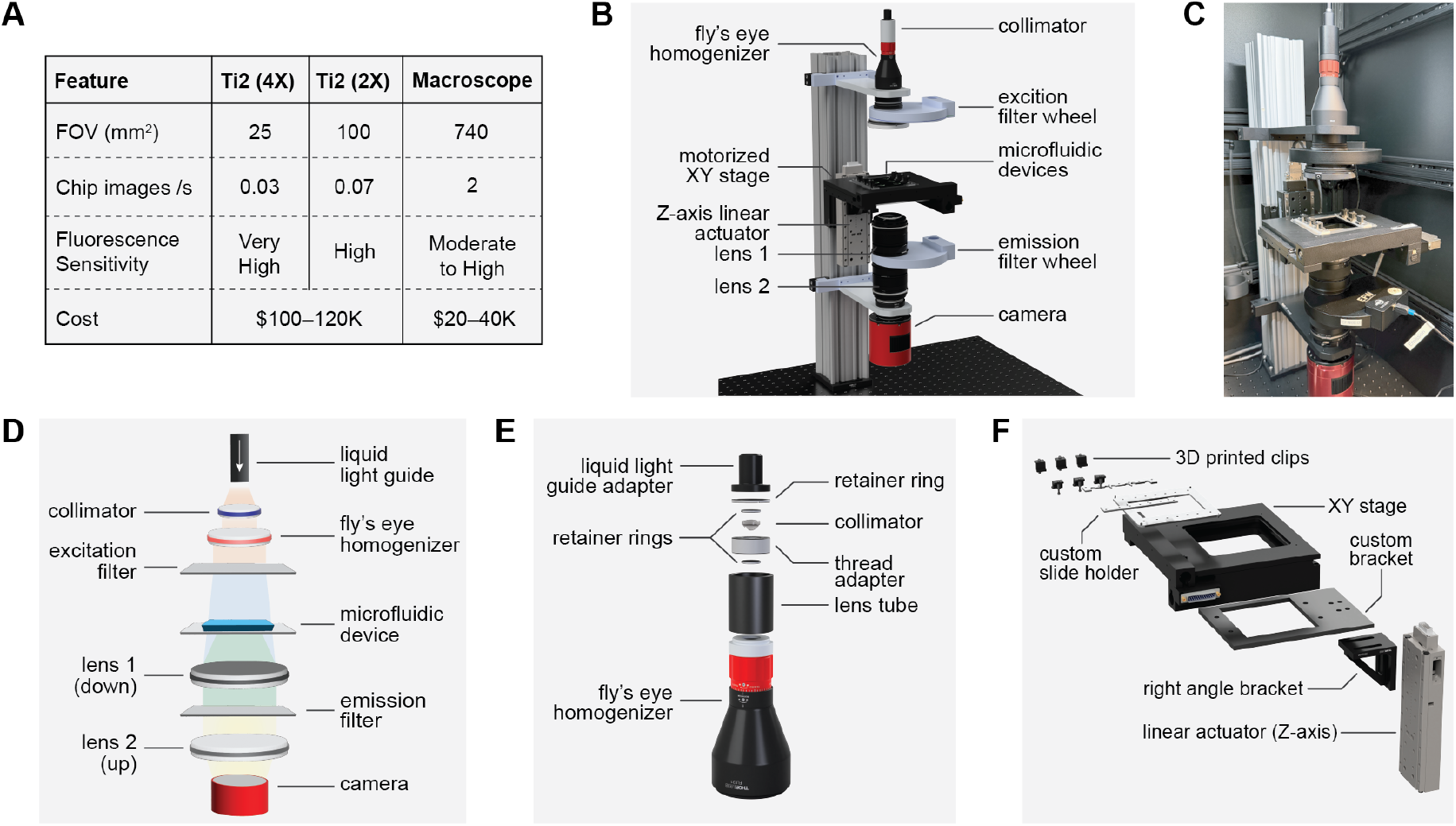
Transfluorescence macroscope for large field of view automated microfluidic imaging. (A) Comparison of key features and cost of commercial Nikon ECLIPSE Ti2 system (equipped with 4X or 2X objectives, as described in Materials and Methods) and custom macroscope described here. (B) Computer-aided design model of the complete macroscope assembly on an optical construction rail mounted on an optical breadboard. (C) Photograph of the completed macroscope optomechanical system within a dark enclosure. (D) Summary of the macroscope’s light path. Diverging output from a liquid light guide is collimated and passed through a fly’s eye homogenizer to achieve wide and flat illumination that is filtered through an OD6 bandpass filter. Light transmitted through or emitted from the sample passes through emission filters mounted in the “infinity space” between the two macro lenses before being re-focused on the sensor. (E) Exploded view of collimation assembly within a lens tube. (F) Exploded view of motorized XYZ stage assembly and custom stage insert consisting of custom laser-cut and 3D-printed components.

As many microfluidic devices are optically transparent (including the polydimethysiloxane (PDMS) devices used for HT-MEK and *k*-STAMMP), we used a transillumination imaging configuration that places the sample between the excitation light source and the tandem objective lens pair (Figs. 1B–D). Unlike traditional epi-illumination tandem lens imaging setups, which require a large gap between lenses to accommodate a filter cube (excitation filter, emission filter, and dichroic mirror) for fluorescence imaging, this trans-illumination configuration requires inserting only an emission filter between the tandem lenses for fluorescence imaging. This allows the tandem lenses to be positioned closer together, significantly reducing image clipping and proportionally increasing the FOV.

To make optimal use of this large FOV, we designed an optical system capable of uniformly illuminating a large area. Here, we collimate and expand light emitted by a Lumencor SPECTRA light engine by routing it through a liquid light guide followed by a fly’s eye homogenizer (a commercial microlens array system that produces a rectangular flat-top beam intensity profile from an input Gaussian beam) (Fig. 1E). A motorized excitation filter wheel positioned directly after the fly’s eye homogenizer allows automated selection of desired excitation wavelengths before reaching the sample (Fig. 1B). The use of a light engine enables flexible excitation across multiple wavelengths; however, this approach is compatible with a wide variety of solid-state light sources.

After reaching the sample, transmitted (bright-field) or emitted (fluorescence) light is propagated via a symmetric stacked tandem lens system composed of two commercial fast (F1.4) prime lenses with manual focus and aperture control. The object-space resolution depends on the ratio of the sensor pixel size (2–7 μm for most commercial CMOS camera sensors) to the primary magnification (1X magnification for identical tandem lenses), corresponding to an expected camera resolution of approximately 5–16 *μ*m where 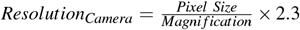 (where 2.3 is a commonly-used sampling rate threshold under the Nyquist-Shannon sampling theorem). ^12^ Thus, this system maximizes FOV while retaining sufficient resolution to observe low-micron-scale features.

Sensitive fluorescence detection in a transillumination configuration requires filters capable of stringently rejecting excitation photons being routed directly towards the camera. Here, we filtered unwanted excitation light by stacking two OD6 bandpass emission filters within custom holders in each slot of the emission filter wheel sandwiched between the lenses of the objective system (Fig. S1). We acquired images using a full-frame camera designed for astrophotography since this class of camera uses large, high-resolution, and actively cooled sensors at substantially lower cost than large-sensor scientific sCMOS cameras. Specifically, we used an ASI 6200MM-Pro camera with a Sony IMX455 CMOS sensor (a back-illuminated, full-frame, 8K-resolution (9576×6388 pixels, 61 megapixels) actively-cooled monochrome sensor with 3.76 *μ*m pixel size). While the full-sensor readout speed of 0.5 s (2 FPS) for the ASI camera is slower than the 10–100 FPS offered by many scientific sCMOS cameras, the ability to record data from across an entire device in a single image leads to a dramatic in-crease in effective temporal resolution.

To automate imaging of multiple samples at once and enable image-based autofocus, we included a motorized XYZ stage system composed of an XY stage with 100 mm by 100 mm of travel (ASI MS-2000 Small XY Stage) and a linear Z stage with 50 mm of travel (ASI LS-50) (Fig. 1F). ^13^ This system has high-resolution rotary encoders capable of sub-micron movement precision in each axis and is controlled with an ASI LX-4000 controller via Micro-Manager drivers for automation. ^14^ We also equipped this stage with a custom insert capable of securely holding three 1×3-inch slides and set screws for stage leveling.

To accommodate all required components within a compact footprint (10×17-inch), we mounted them on an optical construction rail within an extruded-aluminum-framed enclosure to create a dark environment (Fig. 1C). This packaging additionally makes it possible to include fluidic control systems within the same enclosure on a 24×36-inch optical breadboard, providing automated fluidic control and large FOV imaging of multiple devices at a fraction of the cost of commercial systems. ^15,16^

To characterize the optical performance of our setup for brightfield microscopy, we imaged a standard positive USAF-1951 test target (Fig. 2A). As expected, pixels at the edges of the sensor exhibited signs of clipping, with little transmitted light. Nevertheless, propagated sample images covered a substantial portion of the sensor to yield an effective FOV of 34 mm in diameter (~740 mm^2^). To assess bright-field resolution and how it changes across the FOV, we acquired images of the target positioned at the center and far edge of the center (Figs. 2A and S2). When positioned near the sensor center, the vertical and horizontal bars of group 7, element 2 were resolvable, corresponding to an apparent object-space resolution of ~3.5 *μ*m. This resolution is similar to the camera’s pixel size of 3.76 *μ*m, consistent with predictions. When positioned at the sensor edge, the minimum resolvable feature size increased to 3.9 *μ*m (corresponding to element 1 of group 7) (Fig. S2). These images suggest that the macroscope’s bright-field imaging resolution is primarily limited by the sensor’s pixel size, with additional considerations depending on the sample features to be imaged and application-specific performance goals including contrast. ^12^

**Fig. 2:**
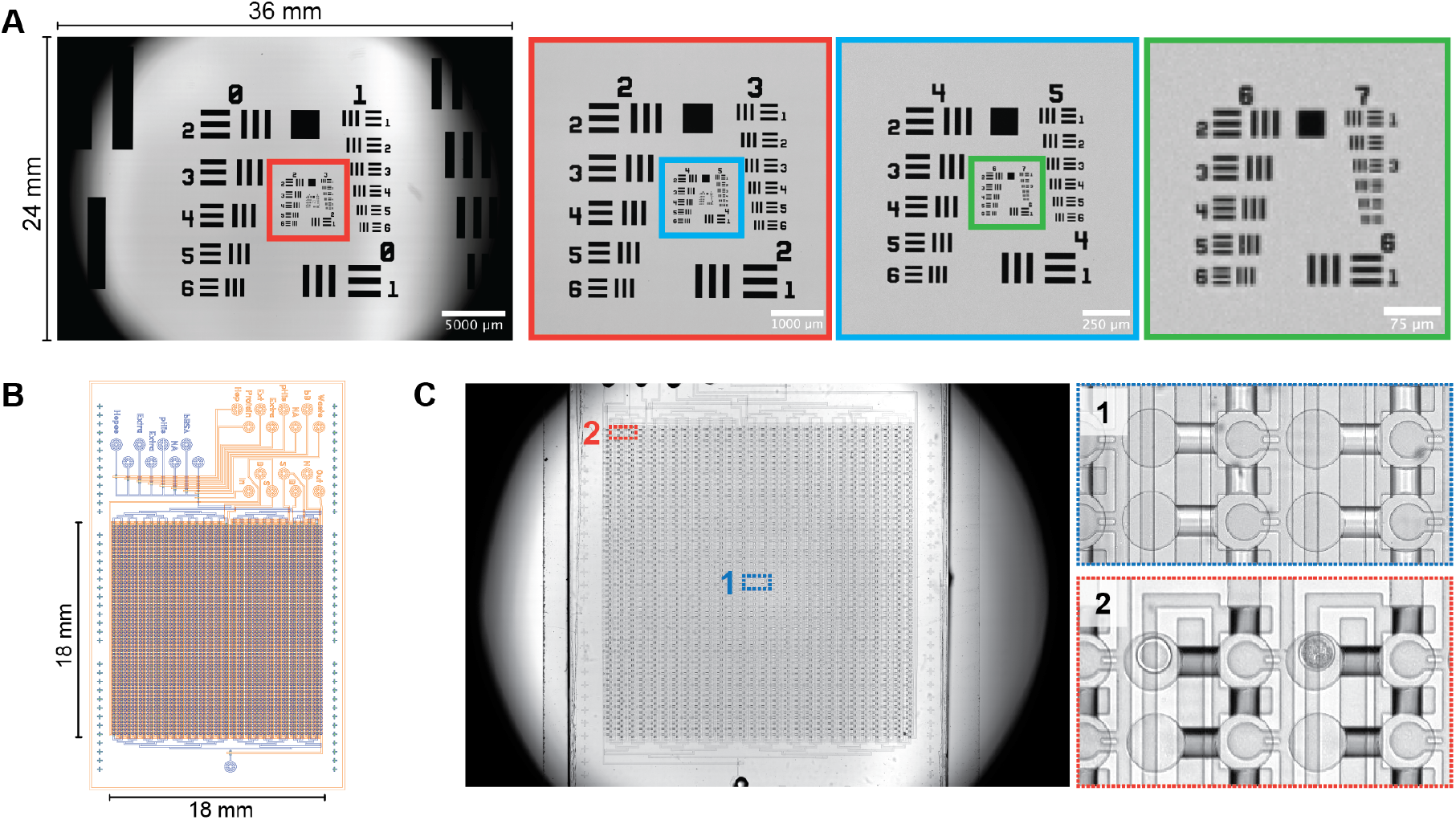
Characterizing macroscope bright-field imaging performance. (A) Bright-field image of a standard USAF test target with expanded inset colored squares encompassing successively higher group numbers (smaller features). (B) Schematic of the HT-MEK microfluidic device showing “flow” (blue) and “control” (orange) PDMS layer features used for on-chip protein expression, purification, and biophysical assays. The ROI is an 18×18 mm square area consisting of circular chambers, flow paths, and overlying integrated valves. (C) Bright-field device image with zoomed-in views of the center (1,blue) and corner (2, red) of the ROI.

To assess the practical utility of this system for bright-field imaging of microfluidic devices, we imaged an HT-MEK microfluidic device with 1,568 225-*μ*m-diameter reaction chambers distributed across an 18 mm × 18 mm area and a minimum feature size of 25 *μ*m (Fig. 2B). ^5^ The plane of focus across the FOV remained flat enough to resolve all key device features, with only a slight loss of focus and soft vignetting at the edge of the ROI (Fig. 2C).

Next, we assessed the sensitivity of fluorescence imaging in different channels under conditions that mimic typical device operation and compared performance to a high-end commercially available microscope system. Specifically, we imaged HT-MEK microfluidic devices loaded with serial dilutions of three common fluorophores (fluorescein, GFP channel; Cy5 (Cyanine-5), Cy5 channel; 6,8-difluoro-7-hydroxy-4-methylcoumarin (DiFMU), DAPI channel) using both our tandem lens transillumination macroscope and a Nikon ECLIPSE Ti2-series commercial epifluorescence microscope equipped with a 1X Plan Achro (NA=0.04), 2X Plan Apo (NA=0.1), and a 4X Plan Apo (NA=0.2) objectives. Specifically, the Ti2-series microscope included a high-powered solid-state light source (Lumencor SOLA U-N Light Engine) for fluorophore excitation and a scientific sCMOS camera (Photometrics Kinetix) with a large sensor (21×21 mm), high resolution (3200×3200 pixels, 10.2 megapixels), high sensitivity (95% peak quantum efficiency), and a pixel size of 6.5 *μ*m. Each fluorophore dilution series was comprised of dye concentrations from 20 nM to 62.5 *μ*M in 1X PBS (pH 7.4); dilutions were imaged in GFP, Cy5, and DAPI channels with exposure times of 50, 50, and 500 ms, respectively. Each image was flat-field corrected using reference images of concentrated fluorophore trapped between two slides (Fig. 3A); after flat-field correction, at each concentration, we calculated the median intensity within each chamber (Figs. 3B and 3C). ^17^

**Fig. 3:**
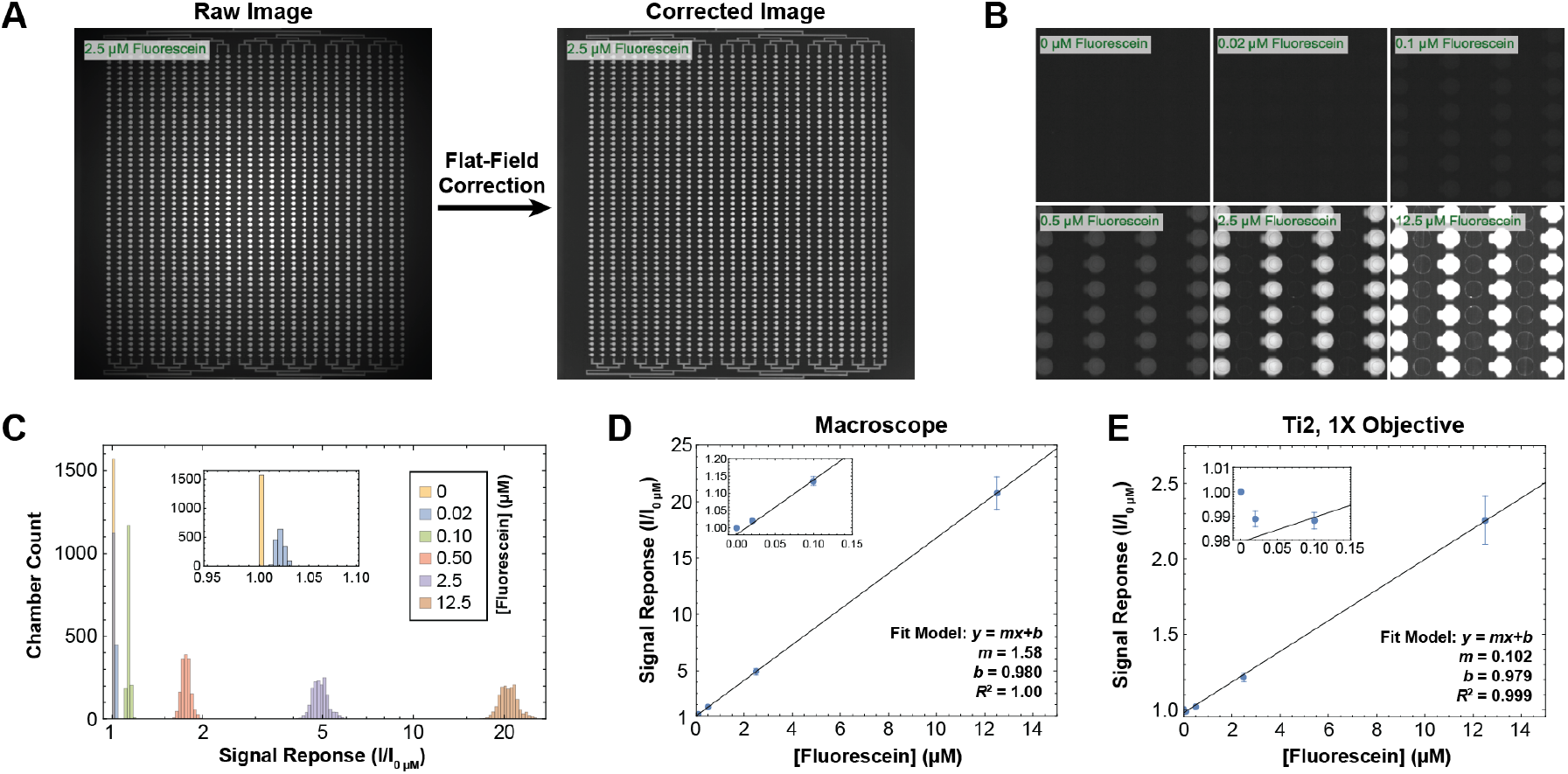
Characterizing macroscope fluorescence imaging sensitivity. (A) Example flat-field correction of a one-shot image of an HT-MEK device containing 2.5 μM fluorescein in PBS (50 ms exposure, GFP channel) using the macroscope. (B) Representative images of a titration series of fluorescein in PBS within central region of the HT-MEK device. Min/Max chamber intensities are set to 0/2000 RFU across all images to improve visual clarity at lower concentrations such that chamber pixel intensities appear saturated at 12.5 μM. (C) Binned histogram of median chamber intensities of fluorescein standard series acquired with the macroscope shown in (B) and normalized to background intensity on a per-chamber basis (1568 chambers, see Materials and Methods). (D) Linear fit to mean of signal response from panel (C) across all chambers. Error bars are ±1 SD from the mean. (E) Linear fit to mean of signal response across all chambers from a fluorescein standard curve acquired on Ti2 comparison system with 1X objective (50 ms exposure, GFP channel). Error bars are ±1 SD from the mean.

For all dyes, fluorescence intensities increased linearly with increasing dye concentration over a wide concentration range (0.1–12.5 *μ*M), as expected. The lowest concentrations detectable above background for the macroscope were <20 nM for fluorescein, <20 nM for Cy5, and <500 nM for DiFMU within device chambers (Figs. 3D and S3). DiFMU is a considerably dimmer fluorophore than either fluorescein or Cy5 (*Brightness* = *ε* × *ϕ*; DiFMU (pH 10): *ε* × *ϕ* = 1.6 × 10^4^ M^−1^cm^−1^, Fluorescein (pH 9): *ε* × *ϕ* = 8.6 × 10^4^ M^−1^cm^−1^, Cy5 (PBS): *ε* × *ϕ* = 6.8 × 10^4^ M^−1^cm^−1^), consistent with the higher limit of detection observed. ^18,19^ Imaging on the Ti2-based setup using the 1X objective yielded a 2–10-fold lower apparent signal response compared with the macroscope across all channels (Figs. 3E and S4–S6). As expected, imaging with either the 2X or 4X objective substantially increased the apparent signal response for all dyes (Figs. S4–S6), but also required tiling images (typically 4–6 and 16–25 tiles per ROI for 2X and 4X objectives, respectively).

To demonstrate an application of this macroscope, we proceeded to quantify enzyme kinetics for a highly proficient (fast) enzyme, the alkaline phosphatase superfamily member PafA, as a model system within the HT-MEK device. ^5^ In this assay, C-terminally mEGFP-tagged enzyme is immobilized on surfaces within >1,500 ~1-nL valved chambers via recruitment to anti-eGFP antibody patterned surfaces; pneumatically-actuated valves within each chamber reversibly protect surface-immobilized enzymes from the surrounding solution to start and stop reactions. Here, we introduced the fluorogenic phosphomonoester substrate 6,8-difluoro-4-methylumbelliferyl phosphate (DiFMUP) into all chambers before raising valves to start the reaction. Once started, surface-immobilized WT PafA hydrolyzed the substrate to produce a fluorescent 6,8-difluoro-4-methylumbelliferone (DiFMU) product, making it possible to follow enzymatic turnover across all chambers in parallel via fluorescence imaging (Fig. 4A). ^5^ Under the conditions of the assay (WT PafA: *k*_cat_/*K*_M_ = 1.2 × 10^6^ M^−1^s^−1^, *k*_cat_ = 120 s^−1^, *K*_M_ =102 *μ*M, ~50 nM in enzyme concentration, and 50 *μ*M substrate), we expect these reactions to reach 50% completion in ~16 seconds, and that obtaining accurate initial rates (corresponding to <20% of full product turnover) would require linear fits of the data within the first ~5 seconds of these progress curves.

**Fig. 4:**
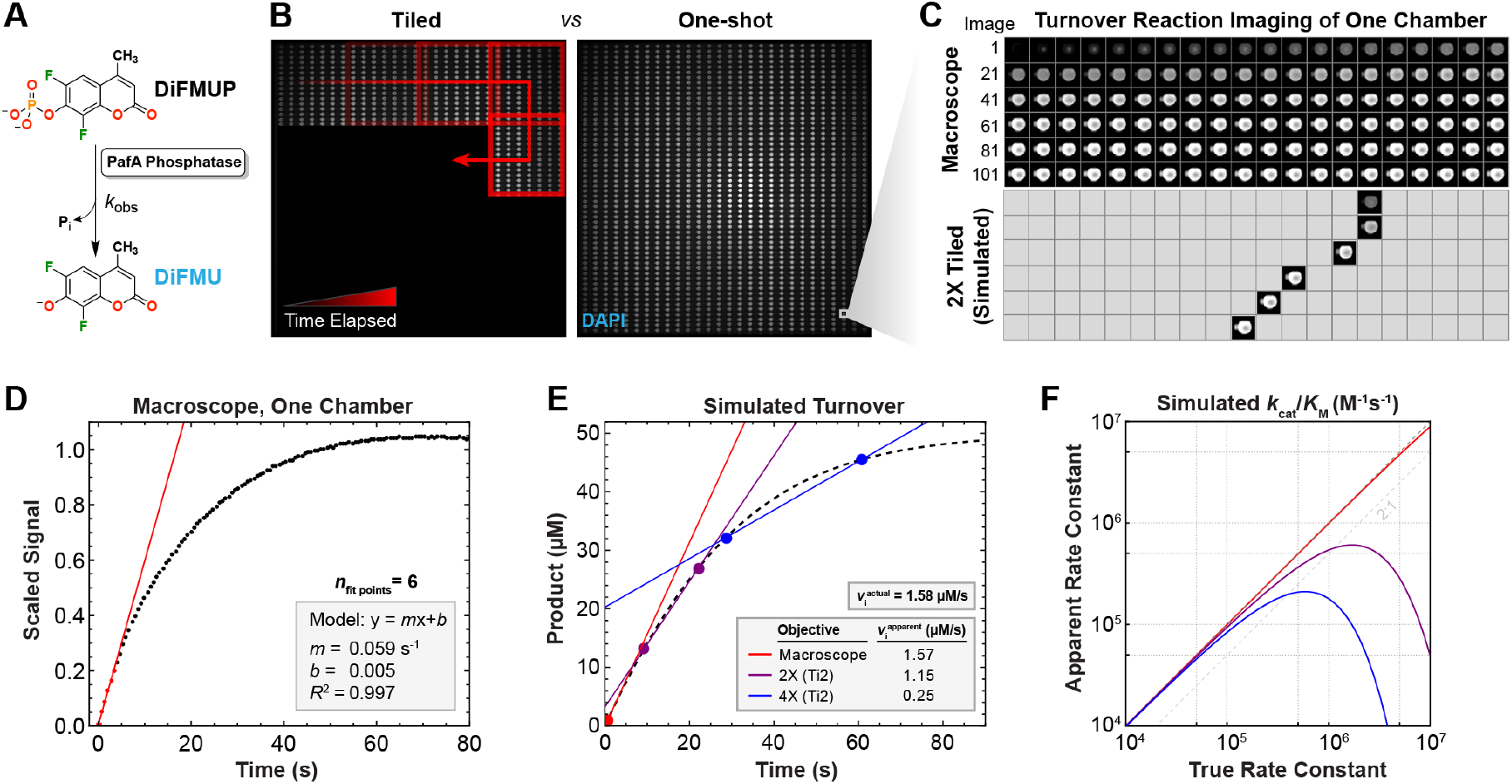
Resolving a rapid enzyme-catalyzed reaction within the HT-MEK device using the macroscope. (A) DiFMUP is hydrolyzed by PafA phosphatase to produce a fluorescent product (DiFMU) and inorganic phosphate. (B) Comparison of tiled imaging with a Ti2-based system using a 4X objective (simulated representation of tiled image acquisition, left) vs one-shot imaging with the macroscope (real image, right) of an HT-MEK chip during an enzyme fluorescence assay kinetic time course. (C) Rapid turnover time-course imaging of PafA-catalyzed DiFMUP (at 50 μM) hydrolysis (top) vs images that would be observed using slower tiled imaging on a Ti2-based system equipped with a 2X objective (bottom) for one example chamber within an HT-MEK chip. (D) Scaled median signal over time within the chamber images shown in the top sub-panel of (C). The initial 6 data points, measured within the first 5 seconds of turnover, are fit to a linear model to yield the apparent initial rate (*m*). (E) A simulated reaction progress curve (black dashed curve) for a PafA-catalyzed reaction with 2-point initial rates fits from observations of a hypothetical chamber near the bottom of the chip using the 2X and 4X objectives on a Ti2-based system vs macroscope. This simulation used a closed-form solution of the integrated Michaelis-Menten equation with kinetic constants for WT PafA with an analogous substrate (*k*_cat_ = 120 ^-1^, *K* _M_ = 102 μM, [E]_0_ = 40 nM, [S]_0_ = 50 μM). ^5^ (F) Simulations of true vs apparent rate constants from 2-point fit initial rates, derived from imaging with the same three objectives and hypothetical chamber shown in panel (E) for enzymes of varying *k*_cat_/*K* _M_. For this simulation, a hypothetical reaction under sub-saturating conditions was modeled as a simple second-order kinetic process (*v*_i_ =(*k*_cat_*/K*_M_)× [*E*]_0_ × [*S*]_0_) with a fixed *K* _M_ (*k*_cat_ : variable, *K* _M_ = 102 μM, [E]_0_ = 40 nM, [S]_0_ = 10 μM).

Previously reported HT-MEK assays profiling PafA employed tiled imaging using a commercial Nikon ECLIPSE Ti-series microscope with a 4X objective such that the effective imaging rate for each chamber was <0.03 Hz (>30 seconds between iterative images) (Fig. 4B). By contrast, the macroscope captured images of the entire device at ~1.7 Hz (~600 ms between iterative images), a >50-fold increase in temporal resolution, enabling accurate capture of initial linear rates (Figs. 4B–D). Even compared to simulations of imaging using a 2X objective on a Ti2 system, the macroscope remained ~20-fold faster (Figs. 4C and 4E).

This increase in imaging speed led to more accurate estimates of fast turnover rates. While the macroscope accurately captured the true initial rate (<1% delta from 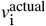) under the conditions of this simulated enzyme turnover assay, simulated slower imaging with a Ti2 and a 2X or 4X objective led to underestimation of the true initial rate of hydrolysis by 1.4-fold and 6.3-fold, respectively (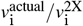 and 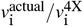, respectively). For *k*_cat_/*K*_M_ values above 10^6^ M^−1^s^−1^, tiled imaging leads to dramatic underestimation of true rates while macroscope imaging remains accurate (Fig. 4F).

## Conclusions

Here, we described a low-cost, multi-channel, 1X transfluorescence macroscope capable of extremely wide-FOV imaging with motorized XYZ stage and filter wheels. This imaging system produces a 34 mm diameter image circle, resolves features in bright-field at the sensor resolution limit (3.8 *μ*m), and quantifies common fluorophores with low-to mid-nanomolar sensitivity within PDMS microfluidic devices. Moreover, this apparatus costs a fraction of the cost of turn-key epifluorescence systems. To demonstrate the macroscope’s capabilities, we used it to accurately quantify fast kinetic turnover of a fluorogenic substrate by PafA phosphatase. These capabilities makes it possible to accurately quantify turnover rates even for enzymes with *k*_cat_/*K*_M_ near the diffusion limit (10^6^−10^8^ M^−1^s^−1^). We anticipate that this setup will prove broadly useful for parallel microfluidic measurements of fast kinetic processes, including fast turnover kinetics (high *k*_cat_) and fast dissociation and association kinetics without a need for the iterative mechanical trapping used by MITOMI-type devices. ^3^ Beyond biochemical applications, this system could also dramatically increase the time resolution of organoid and sub-mm organism imaging over large ROIs. ^20,21^

For more demanding applications, fluorescence sensitivity and temporal resolution could be increased via incorporation of higher-end scientific cameras and inclusion of additional filters to more effectively block excitation light. The camera we used has a minimum read-out time of 0.5 s, defining the maximum theoretical sampling rate to 2 Hz. However, a higher-end camera with faster readout could increase the maximum imaging rate to ~20 Hz, assuming use of a camera capable of 120 FPS (sensor readout rate) and exposure time of 40 ms. While clipping by internal elements of the tandem lenses used here precludes imaging even larger FOVs, custom-designed optics could increase FOV by filling a full-frame or medium-format sensor with the image circle. These custom optics could be additionally designed to reduce curvature of the focal plane and optimize depth of field for enhanced imaging uniformity.

Here, we provided a stringent benchmark of macroscope performance by comparing images to those acquired using a state-of-the-art comparison system (Nikon Ti2) equipped with large-diameter filters (32 mm), a camera with very large sensor (433 mm^2^), and high-performance 2X and 4X high-NA objectives previously selected to maximize the FOV. Compared to many commercial systems, we expect that the macroscope would yield even larger improvements to sensitivity, FOV, and temporal resolution.

## Author Contributions

D.A.M. conceived of and co-supervised the project. P.M.F. cosupervised the project and acquired funding. D.A.M and A.L. designed experiments and collected and analyzed data. D.A.M, A.L., and P.M.F. wrote and revised the manuscript.

## Acknowledgments

This work was partially supported by an NIH Director’s Pioneer Award (DP1CA290563) awarded to P.M.F, and P.M.F. is a Chan Zuckerberg Biohub investigator. We thank Youngbin Lim for his technical assistance and helpful discussion.

## Data availability

Data supporting the findings of this study are available within the article or at an Open Science Foundation (OSF) repository at https://doi.org/10.17605/OSF.IO/DY5XP. OSF repository data include computer-aided design models of custom-manufactured parts described herein.

