## Supplementary Information for "Large field of view fluorescence imaging of microfluidic devices with a tandem-lens macroscope^†^"

**Figures S1 to S6** **9**

**Table S1** **15**

**References** **16**

### Materials and Methods

#### Macroscope imaging system construction

##### Design rationale and build summary

A macroscope build list of components and their prices (as of early 2024) is provided in Table S1. For second-hand components, approximate prices based on listings as of early 2024 are given. The build description below generally references parts by name, and specific part numbers, manufacturers, and vendors can be found in the build list.

**Motion control.** The macroscope utilizes a motorized stage system providing high-precision (sub-micron) XYZ motion control with high-quality linear encoders with build inspired from an ASI manufacturer manual for the ASI LS-Series linear positioning stages (*1*). This type of motion control system would be expected to cost in excess of $15,000 USD when new, but high-precision second-hand stages can be sourced at significantly lower cost. Thus, we built a complete XYZ motion control system from high-accuracy second-hand ASI components, including a motion controller with serial computer interface (ASI LX-4000). This build can routinely be achieved from the second hand market at a total cost of approximately $2,500 USD. This motion control system enables long range of travel ($100 \times 100 \times 50$ mm) and highly-precise (sub-micron repeatability) XYZ control for up to three $1 \times 3$ inch slides, and thus three HT-MEK microfluidic chips, simultaneously.

**Excitation light treatment.** The macroscope's optical train utilizes a standard liquid light guide to route output from an incoherent, high-powered, LED light source. In our build, we used a state-of-the-art Lumencor SPECTRA III light engine since this light source was on hand and offered bright, multi-channel illumination perfect for optical system development. This specific light engine is not necessary to complete this build, and multi-wavelength light sources with similar core features in fewer channels (e.g., Thorlabs CHROLIS 6-channel source or Thorlabs LED4D 4-Wavelength source) can be purchased at substantially lower cost (approximately $12,500 USD and $6,700 USD, respectively, as of Fall 2025). A liquid light guide enabled interfacing with a wide variety of light sources without altering mounts and provided flexibility in system packaging (placing the source outside of an enclosure, for example). Transmitted light is collimated at the top of the macroscope optical train, then expanded with a commercial fly's eye homogenizer to create a rectangular output with approximately top-hat intensity profile spanning the large image circle of the lens system. This approach was found to be superior to a simple Galilean beam expander, providing a significantly more uniform beam profile and higher irradiance at the sample plane.

The shaped excitation beam is then filtered through a 50-mm-diameter OD6 bandpass excitation filter (fluorescence bandpass filters used were from Chroma Technology Corp.) mounted in a low-profile electronic filter wheel (ZWO Astro) with 5 filter slots, equipped with DAPI, GFP, and Cy5-compatible excitation filters in our default configuration. Filtered excitation light is aligned with the central axis of the lens system below and positioned to approximately focus the homogenizer output at the imaging front focal plane. To achieve bright-field imaging on the macroscope, we used multi-channel excitation in red, greed, and blue channels simultaneously at very low power. We attenuated the excitation light beam with an OD 2.0 absorptive ND filter (Thorlabs) loaded in the excitation filter wheel, and used an empty emission filter cell to pass all transmitted light to the camera sensor.

**Emission light treatment.** Emitted light from the imaging front focal plane is collected via a finite-corrected optical system utilizing a face-to-face, tandem-lens macro configuration related to previously described systems (*2*). Our system differs from most other reported tandem-lens macro configurations in that we use a symmetric configuration with the same lens on both sides. This symmetric configuration produces an approximately 1:1 image (1X magnification) of the sample on the camera sensor, and was selected to produce a very large image circle. In this type of system, each lens is set at infinity focus, them mechanically coupled face to face such that the "sensor side" of each lens is sitting outward. This configuration can produce extremely shallow (down to low micron) depth of field and produce a sharp image across a very large (10s of mm) field of view.

To build our lens system, we considered fast, full-frame commercial lenses with manual aperture and focus control, and tested several commercial lenses meeting these criteria to rank them based on image circle diameter, vignetting, aberration, relative numerical aperture, and other factors. The degree of hard vignetting, and thus final image circle seen on the sensor, soft vignetting, and spherical aberration, were found to vary widely lens to lens due to differences in internal lens architecture and face diameter. We settled on a low cost, full-frame-compatible Rokinon 85 mm F1.4 lens that yielded a good tradeoff of cost, compactness, apparent numerical aperture, and front face diameter (which was an important constraint for emission filter wheel mounting). We noted in our testing that extremely fast lenses (i.e., F1.2 or less) produced depth of fields so shallow that field curvature (Petzval curvature) became very noticeable, and it also became challenging to reliably align the stage sample plane with the focal plane. Incorporating additional axes of precise stage motion adjustment could make these faster, and thus higher numerical aperture, lens systems more appealing for use in our macroscope system from the perspective of aligning sample and lens focal planes, but custom optics would be required to reduce the field curvature.

Emission filtering was performed within the approximately infinity-corrected space between the tandem lenses to minimize distortion. Since hard vignetting was seen to worsen (image circle reduces) as a function of tandem lens separation, we minimized separation of the lenses by directly mounting a second 5-position low-profile electronic filter wheel to the front faces of (between) the tandem lenses. Since excitation light shines directly into the lens system, we stacked two (2) identical OD6 bandpass emission filters face-to-face within the filter wheel using a custom filter cell and spacer (Figure S1).

#### Choice of a transillumination optical system

By utilizing a trans-illumination configuration, our system can maintain a larger field of view than would have been possible by using epi-illumination. This is because with a dichroic positioned between the tandem lenses, lenses would have to be separated by a much larger distance, which would increase the degree of hard vignetting (clipping) and thus reduce the diameter of the image circle produced on the sensor. In addition, although including a second emission filter per channel to adequately block excitation light in our trans-illumination system incurs additional costs, our approach offsets that cost by dispensing with a dichroic mirror and mounts (cubes) required to co-align paired filters and a dichroic.

##### Choice of imaging camera

We selected a camera designed for astrophotography (ZWO, ASI6200MM Pro) to cost-effectively meet the needs of our imaging application. This camera, and other similar astrophotography cameras, offer very large (i.e., full-frame, 8K-resolution) CMOS based sensors with 16-bit analog-to-digital converters and active cooling with high peak quantum efficiency (>90%), while offering favorable maximum frame rates (2 Hz) at modest cost (~$4,000 USD). In comparison, high-performance scientific cameras (e.g., Teledyne Photometrics Kinetix, Teledyne Vision Solutions) for several times the price offer some similar features but offer very high speed readout and data transfer rates (>80 FPS full-frame readout in high-dynamic range mode) that are often not required for microfluidic applications. In addition, most possess significantly smaller and lower resolution sensors (the majority of scientific sCMOS sensors on the market are <3200×3200 pixels).

##### Software control

The light source, excitation filter wheel, XYZ stage, emission filter wheel, and camera are controlled and synchronized using the Micro-Manager open source microscope control software package (*3*). By using Micro-Manager and configuration presets, this macroscope imaging system was easily interfaced with an existing, previously-described Python package capable of scripted and coordinated microfludic valve control and imaging (*4*). By abstracting hardware-level configuration and communication behind Micro-Manager presets and API calls, our macroscope can be easily upgraded and extended with additional commercial components without requiring significant changes to custom automation code. Additionally, since high-intensity unfiltered excitation light focused on the sensor could damage the camera, use of Micro-Manager presets that match the excitation and emission filters with corresponding light source channel and appropriate output power prevents accidental system damage.

#### Ti2-based commercial epifluorescence microscope comparison system

As a comparison imaging system for benchmarking, we used a commercial, state-of-the-art, motorized Nikon ECLIPSE Ti2-E inverted epifluorescence microscope system equipped with extremely high-end components. This microscope was equipped with a high-resolution motorized ASI XY stage, motorized focus control (Z-stage), a Lumencor SOLA U-N solid-state white light source, a Teledyne Vision Solutions Photometrics Kinetix camera, and Nikon high signal-to-noise BL Series (large FOV) fluorescence filter cubes (DAPI SOLA, GFP, and Cy5 cubes were used for testing in this work). We used 2X and 4X objectives of the highest numerical aperture readily available (Nikon CFI Plan Apo Lambda D series) and we used the only available Nikon 1X objective (CFI Plan Achro). At the time of purchase, this system had a total cost of over $100,000 USD. If equipped with a Lumencor SPECTRA III light source, as used for the macroscope build, the total build cost of this system would have been approximately $115,000 USD.

#### HT-MEK microfluidic device fabrication

The two-layer PDMS devices (HT-MEK chips) used here were fabricated as previously described from casting of photolithographically-produced silicon wafer molds, also made as previously described (*5*). Device and mold fabrication were performed within class 10,000 and class 1,000 clean rooms, respectively.

#### On-chip fluorescence standard curves

##### Fluorophore stock preparation

Dilution series of fluorescein disodium salt (VWR cat. no. 97062-186), 6,8-difluoro-7-hydroxy-4-methylcoumarin (DiFMU, ThermoFisher Scientific, cat. no. D6566), and Cyanine5 (Cy5, MedChem Express, cat. no. HY-D0821) dyes were prepared in 1X phosphate-buffered saline (PBS, ThermoFisher Scientific, cat. no. 10010023) from concentrated stocks in anhydrous dimethyl sulfoxide (DMSO, ThermoFisher Scientific, cat. no. D12345). Each standard was prepared as a serial dilution at the following concentrations (µM): 0, 0.02, 0.1, 0.5, 2.5, 12.5, and 62.5. For some imaging exposure times, 62.5 µM of fluorophore saturated the camera sensor, in which case that concentration was omitted from downstream analyses.

**Device setup.** Brightfield images were acquired with dry (*i.e.*, no fluidic control lines were connected) HT-MEK chips on both the macroscope and our benchmark Ti2-based imaging system. For fluorophore dilution series imaging, pressurized fluidic control lines containing water were connected to the chips as previously described (*6*).

**Image acquisition.** Imaging and fluidic control of chips for dilution series was scripted using the customized Python-based hardware automation framework described above (Software control) within a Jupyter Notebook. The apparent focal length of the lens system was found to have small dependencies on the fluorescence channel used (emission filter identity) with differences on the order of approximately $\leq$100 µm, and so prior to standard curve acquisition, offsets in imaging system focus Z-positions were measured using a low concentration of each fluorophore and corresponding filter and coded into the standard curve acquisition scripts so that images for each standard were taken at the optimal Z-position corresponding to that channel. A 10% overlap between tiles was used for all images acquired on the benchmarking Ti2 setup using 2X or 4X objectives.

**Flat-field correction and stitching.** Images taken on both the macroscope and Ti2 were first flat-field corrected to remove vignetting-dependent brightness differences across the field of view. Flat-field correction was performed as previously described (*4*). Briefly, flat-field images were acquired on both imaging apparatuses by taking images of concentrated fluorophore used in each standard series in its respective imaging channel. Correction was then applied as previously described (*7*). Flat-field corrected images acquired on the Ti2 equipped with a 2X or 4X objective were then "stitched" to construct a complete chip image using a custom Python package previously reported (*4*). As images acquired on the macroscope and on the Ti2 equipped with the 1X objective contained the entire region of interest, no tiling was required in these cases. Then, images were subjected to feature finding (chamber edge detection) and fluorescence intensity quantification, producing summary reports of apparent fluorescence intensity within each chamber at each concentration point. Image processing was also performed with a custom Python package in a Jupyter Notebook as previously reported (*4*).

#### PafA turnover assay

##### PafA construct design and cloning

A fusion construct of PafA phosphatase (Uniprot ID Q9KJX5) and monomeric eGFP (mEGFP) was designed for *in-vitro* transcription/translation (IVTT) with PURExpress (New England Biolabs). This expression construct was generated from a synthesized gene block (Integrated DNA Technologies) consisting of a methionine start codon followed by codons for residues 21–546 of PafA phosphatase, codon-optimized for *E. coli*, cloned into a backbone vector (PURExpress DHFR Control Plasmid, New England Biolabs, with DHFR coding sequence removed) containing a downstream flexible Ser/Gly linker (amino acid sequence "-GGGSGGGGSG-") and fused C-terminal mEGFP. Cloning was performed via HiFi assembly (New England Biolabs) with linearized vector bearing complementary overhangs, and sequence-verified, miniprepped plasmid was used for downstream IVTT.

##### Device setup and patterning

An HT-MEK chip was pressurized and surface patterned to immobilize a patch of anti-eGFP antibody on the surface within each chamber, as previously reported (*4*).

##### PafA expression, purification, and assay with HT-MEK

**Off-chip cell-free expression and on-chip purification.** PafA-mEGFP fusion was expressed off-chip via *in vitro* transcription-translation (IVTT) with PURExpress reconstituted cell-free expression system (New England Biolabs, cat. no. E6800L), and this expression mixture was then flowed over one-half of a patterned HT-MEK chip to immobilize and purify the expressed enzyme. First, an 18 µL expression reaction was prepared on ice by combining 14 µL of a master mix consisting of 4:3 parts PURExpress components A and B, 1 µL of recombinant RNAse Inhibitor (Promega, cat. no. N2511) at 16 U/µL, 2 µL of plasmid DNA template at ~50 ng/µL, and 1 µL of $ZnCl_2$ at 2 mM in water. This reaction was incubated in a thermocycler for 2 hours at 37 ºC, then 1 hour at 23 ºC to express PafA fusion protein. Then, the IVTT reaction was diluted 5-fold into a buffer consisting of 100 mM MOPS, 500 mM NaCl, 100 µM $ZnCl_2$, and 1 mg/mL BSA at pH 8.0 (1X PafA Reaction Buffer), and this IVTT dilution was flowed over the patterned HT-MEK chip for about 10 min with the button valves in the open state. Following binding, excess unbound enzyme was flushed out of the chip by flowing 1X PafA Reaction Buffer for an additional 10 min with the button valves closed, and bound enzyme was visualized on the macroscope by imaging in the GFP channel.

**SDS wash to remove contaminating enzyme.** The chip was washed with an SDS solution, similarly to previously reported, to remove contaminating PafA non-specifically bound to chamber walls (*4*). This wash denatures and inactivates PafA that is not mechanically protected by (specifically recruited beneath) a valve within each chamber. To do this, a solution of 1% SDS in 1 M HEPES buffer at pH 7.3 was flushed continuously through the chip for 20 minutes with these enzyme-bearing valves closed. Then, the chip was flushed copiously with 1X PafA Reaction Buffer prior to turnover assays.

**On-chip turnover assay with DiFMUP.** A kinetic time-series turnover assay was performed on the chip containing surface-recruited and purified PafA using 6,8-difluoro-4-methylumbelliferyl phosphate (DiFMUP, AAT Bioquest, cat. no. 11627) substrate. DiFMUP is an activated fluorogenic phosphomonoester substrate similar to previously reported coumarin substrates of PafA, and was expected to be rapidly hydrolyzed by PafA (*4*). A concentrated stock solution of DiFMUP in DMSO was used to generate a 50 µM stock of DiFMUP in 1X PafA Reaction Buffer. This DiFMUP solution was flowed through the chip for about 8 minutes with enzyme-bearing valves closed (to avoid cleaving substrate prematurely), and then these valves were opened to expose enzyme and initiate reactions while valves separating adjacent chambers were simultaneously shut. Rapid imaging was then initiated (as described above, Software control) in the DAPI channel with 50 ms exposure times to quantify fluorescent product (DiFMU) generation over time.

##### Turnover image analysis

Image processing of macroscope DAPI-channel images from DiFMUP turnover kinetic timecourses was performed as previously reported to extract median chamber intensities for all images at each timepoint (*4*). These intensities were background intensity (intensity at $t$ = 0 s after reaction initiation) subtracted and normalized to the final intensity (at complete product turnover) to generate relative turnover curves used to assess accuracy of initial rate fitting across each chamber. Linear least-squares fitting was performed to estimate initial rates.

### Figures S1 to S6

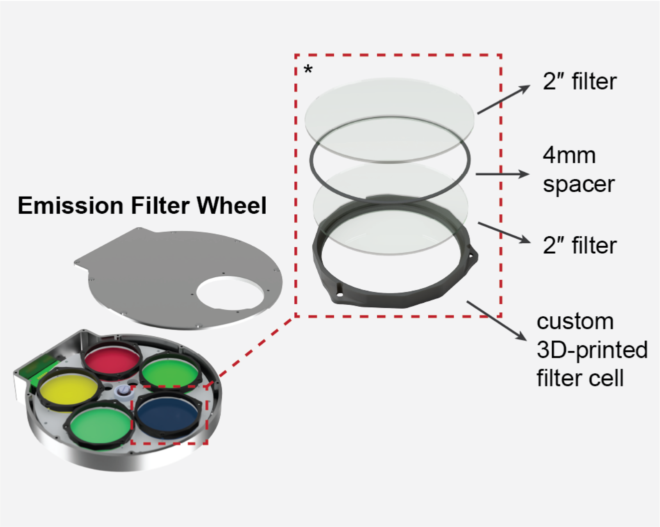

**Fig. S1.** Motorized emission filter wheel assembly containing two stacked OD6 bandpass filters housed in a custom filter cell for each channel.

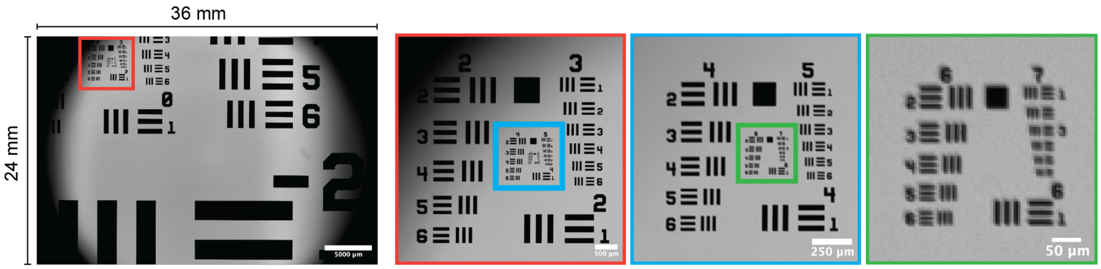

**Fig. S2.** Bright-field image of USAF-1951 test target centered near edge of the macroscope's filed of view.

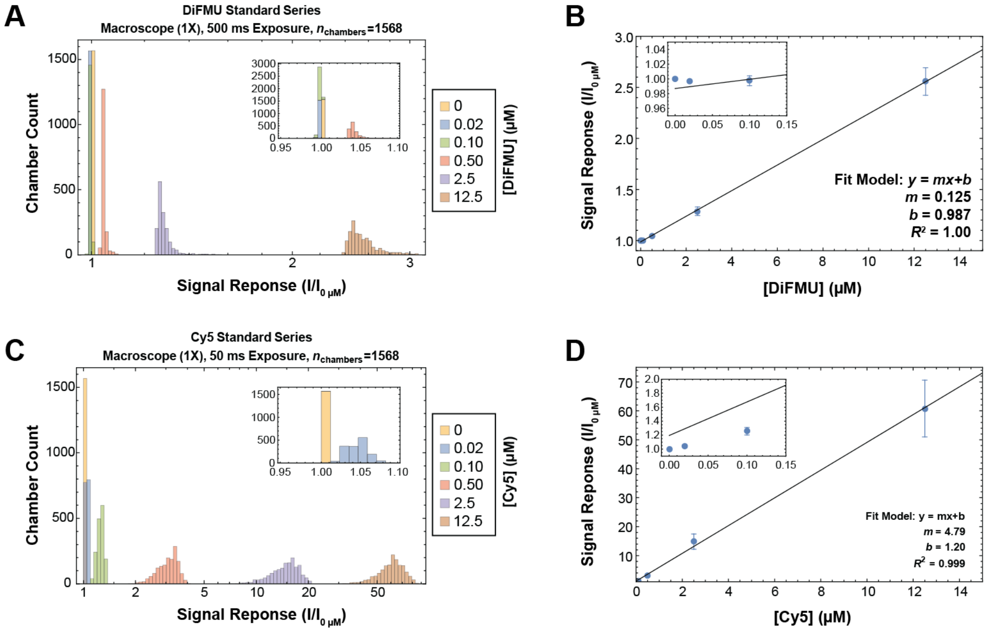

**Fig. S3.** DAPI and Cy5 standard curves on an HT-MEK device acquired with the macroscope. (A & C) Binned histogram of median chamber intensities of DiFMU and Cy5 standard series, respectively, normalized to background intensity on a per-chamber basis. (B & D) Linear fit to mean of signal response from panels (A) & (C), respectively, across all chambers. All error bars are ±1 SD from the mean.

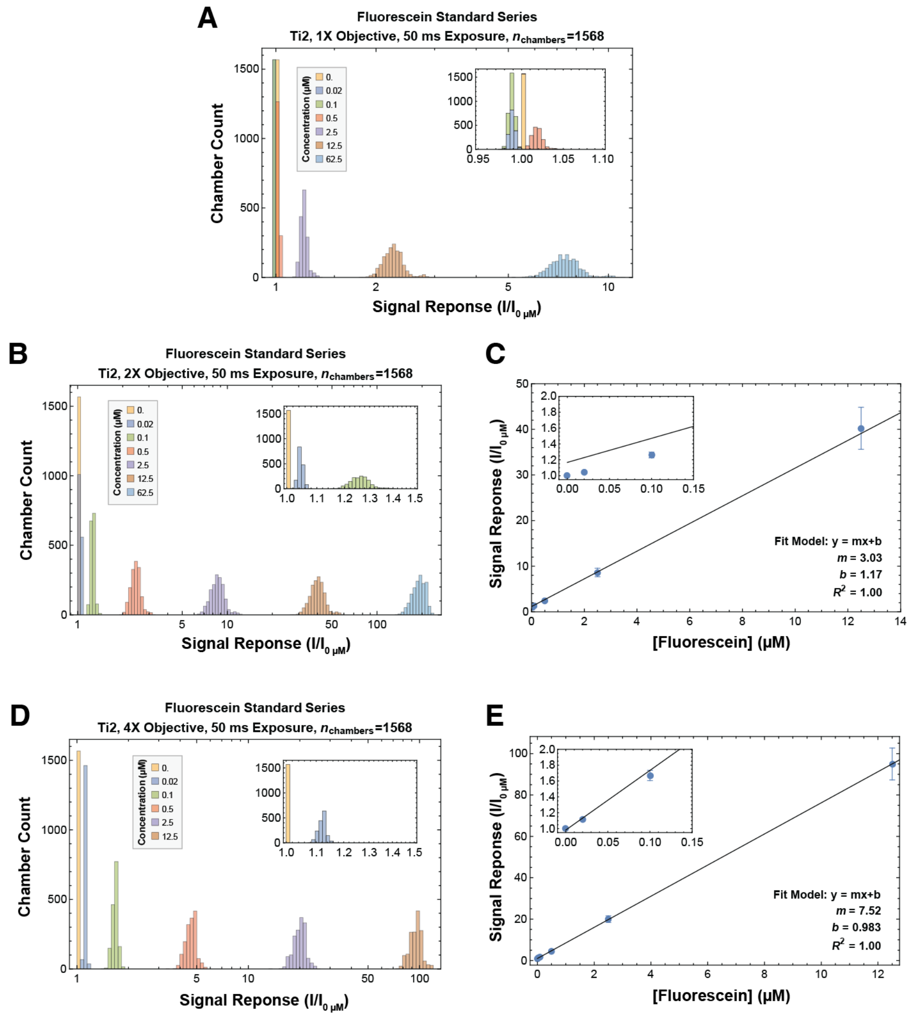

**Fig. S4.** Fluorescein standard curves on an HT-MEK device acquired with a Ti2-based imaging system. (A, B, & D) Binned histogram of median chamber intensities of fluorescein standard series obtained with a 1X, 2X, and 4X objective, respectively, normalized to background intensity on a per-chamber basis. (C & E) Linear fit to mean of signal response from panels (B) & (D), respectively, across all chambers. All error bars are ±1 SD from the mean.

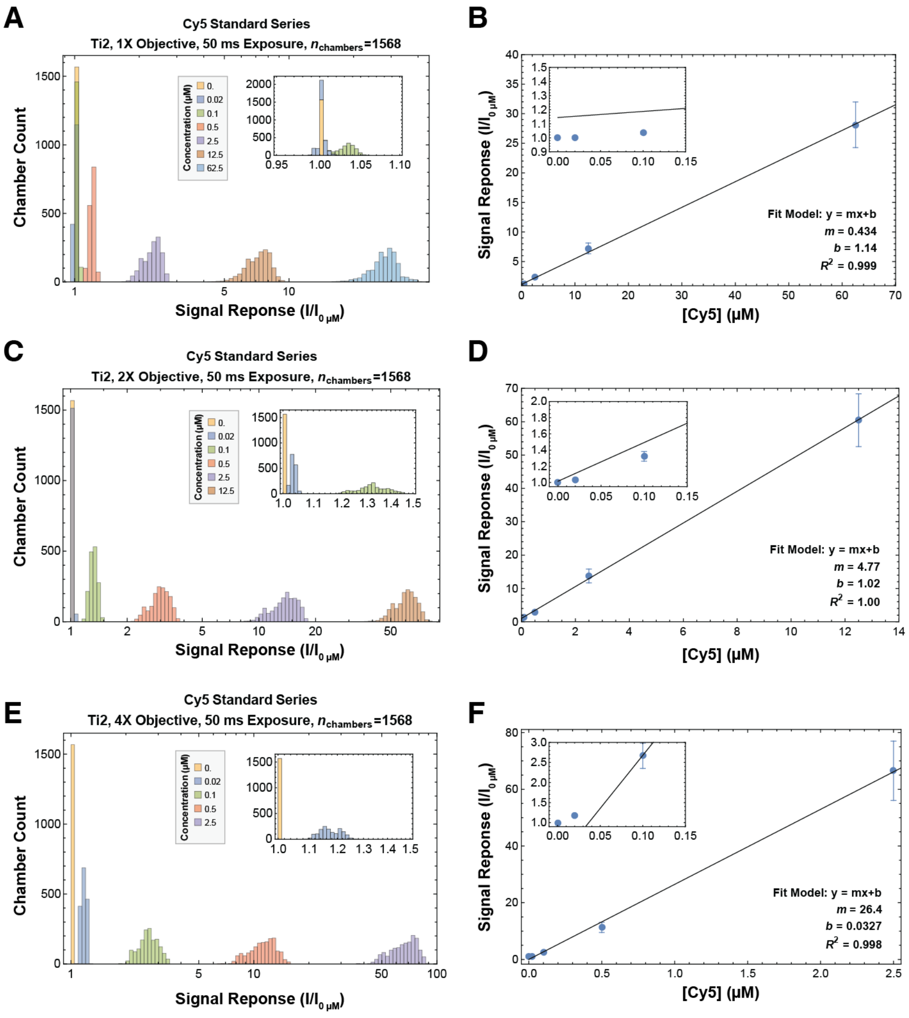

**Fig. S5.** Cy5 standard curves on an HT-MEK device acquired with a Ti2-based imaging system. (A, C, & E) Binned histogram of median chamber intensities of Cy5 standard series obtained with a 1X, 2X, and 4X objective, respectively, normalized to background intensity on a per-chamber basis. (B, D, & F) Linear fit to mean of signal response from panels (A), (C) & (E), respectively, across all chambers. All error bars are ±1 SD from the mean.

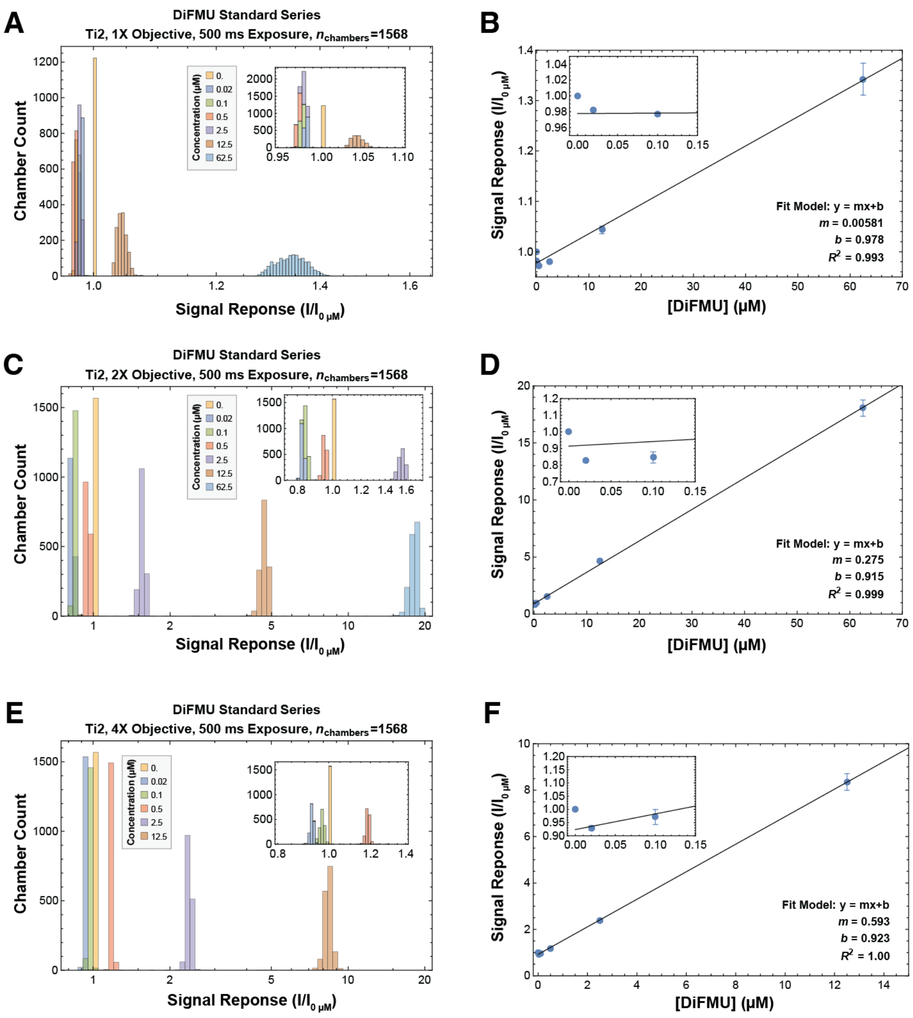

**Fig. S6.** DiFMU standard curves on an HT-MEK device acquired with a Ti2-based imaging system. (A, C, & E) Binned histogram of median chamber intensities of DiFMU standard series obtained with a 1X, 2X, and 4X objective, respectively, normalized to background intensity on a per-chamber basis. (B, D, & F) Linear fit to mean of signal response from panels (A), (C) & (E), respectively, across all chambers. All error bars are ±1 SD from the mean.

### Table S1

| Component | Subcomponent | Manufacturer | Vendor | Part Name | Manufacturer SKU | Count | Unit Price | Total Price | Note |
|---|---|---|---|---|---|---|---|---|---|
| **Camera** | – | ZWO | Agena Astro | ASI6200MM | ASI6200MM-P | 1 | $3,799.00 | **$3,799.00** | |
| **Lens system** | Lens | Rokinon | Amazon | Rokinon Series II 85mm F1.4 Lens | SE85AE-N | 2 | $279.00 | $558.00 | |
| **Lens system** | Coupler | PreciseParts | PreciseParts | Custom ZWO EFW 5x2 filter wheel M54 to generic filter 72 mm | – | 2 | $116.00 | $232.00 | |
| | | | | | | **Component Total** | | **$790.00** | |
| **Stage** | Motion Control | ASI | Ebay | MS-2000 Small XY Stage | MS-2000 | 1 | $1,000.00 | $1,000.00 | Estimated Price |
| **Stage** | Motion Control | ASI | Ebay | LS-50 Linear Stage | LS-50 | 1 | $750.00 | $750.00 | Estimated Price |
| **Stage** | Motion Control | ASI | ASI | LX-4000 Controller | LX-4000 | 1 | $750.00 | $750.00 | Estimated Price |
| **Stage** | Motion Control | Xometry | Xometry | Laser cut stage bracket, aluminum | – | 1 | $57.69 | $57.69 | |
| **Stage** | Stage Insert | Xometry | Xometry | Laser cut stage insert, aluminum | – | 1 | $29.32 | $29.32 | |
| **Stage** | Stage Insert | Xometry | Xometry | Laser cut stage retainers, aluminum | – | 2 | $13.51 | $27.02 | |
| **Stage** | Stage Insert | – | – | 3D-printed slide clamp, ABS plastic | – | 6 | – | – | Printed in house at nominal cost |
| **Stage** | Stage Insert | McMaster-Carr | McMaster-Carr | 18-8 Stainless Steel Socket Head Screw, 2-56 Thread Size, 5/16" Long | 92196A078 | 14 | $0.08 | $1.12 | |
| **Stage** | Stage Insert | McMaster-Carr | McMaster-Carr | 18-8 Stainless Steel Socket Head Screw, 2-56 Thread Size, 5/32" Long | 92196A075 | 6 | $0.09 | $0.54 | |
| **Stage** | Stage Insert | McMaster-Carr | McMaster-Carr | 18-8 Stainless Steel Coupling Nut, 2-56 Thread Size | 90268A203 | 6 | $2.42 | $14.52 | |
| **Stage** | Stage Insert | McMaster-Carr | McMaster-Carr | 18-8 Stainless Steel Nylon-Tip Set Screw, 2-56 Thread Size, 3/16" Long | 90291A847 | 4 | $1.26 | $5.04 | |
| **Stage** | Stage Insert | McMaster-Carr | McMaster-Carr | 18-8 Stainless Steel Narrow Hex Nut, 2-56 Thread Size | 90730A003 | 8 | $0.04 | $0.32 | |
| | | | | | | **Component Total** | | **$2,635.57** | |
| **Light path** | Filter wheel | ZWO | Agena Astro | ZWO EFW (5 x 2") | EFW-5x2 | 2 | $299.00 | $598.00 | |
| **Light path** | Filter wheel | – | – | 3D-printed filter cell, ABS plastic | – | 5 | – | – | Printed in house at nominal cost |
| **Light path** | Filter wheel | – | – | 3D-printed filter cell spacer, ABS plastic | – | 5 | – | – | |
| **Light path** | Excitation Filter | Chroma Technology | Chroma Technology | ET376/30x 50mm Dia Mounted 2.3mm thick ring | ET376/30x | 1 | $865.00 | $865.00 | |
| **Light path** | Excitation Filter | Chroma Technology | Chroma Technology | ET470/40x 50mm Dia Mounted 2.3mm thick ring | IN100494 | 1 | $825.00 | $825.00 | |
| **Light path** | Excitation Filter | Chroma Technology | Chroma Technology | ET620/60x 50mm Dia Mounted 2.3mm thick ring | ET620/60x | 1 | $865.00 | $865.00 | |
| **Light path** | Excitation Filter | Thorlabs | Thorlabs | Unmounted Ø2" Absorptive ND Filter, OD 2.0 | NE2R20B | 1 | $93.18 | $93.18 | |
| **Light path** | Emission Filter | Chroma Technology | Chroma Technology | ET450/50m 50mm Dia Mounted 2.3mm thick ring | ET450/50m | 2 | $865.00 | $1,730.00 | |
| **Light path** | Emission Filter | Chroma Technology | Chroma Technology | ET525/36m 50mm Dia Mounted 2.3mm thick ring | IN044448 | 2 | $825.00 | $1,650.00 | |
| **Light path** | Emission Filter | Chroma Technology | Chroma Technology | ET690/50m 50mm Dia Mounted 2.3mm thick ring | ET690/50m | 2 | $865.00 | $1,730.00 | |
| | | | | | | **Component Total** | | **$8,356.18** | |
| **Light path** | Collimator | Thorlabs | Thorlabs | Ø3 mm LLG to SM1 Adapter | AD3LLG | 1 | $38.32 | $38.32 | |
| **Light path** | Collimator | Thorlabs | Thorlabs | SM1 Lens Tube Sleeve 1.5" Long | SM1M15 | 1 | $19.84 | $19.84 | |
| **Light path** | Collimator | Thorlabs | Thorlabs | Thick SM05 Int. to SM1 Ext. Adapter | SM1A6T | 1 | $23.41 | $23.41 | |
| **Light path** | Collimator | Thorlabs | Thorlabs | 1/2" retaining ring | SM05RR | 1 | $4.33 | $4.33 | |
| **Light path** | Collimator | Thorlabs | Thorlabs | Ø12.7mm, F=8mm, Aspherical Condenser lens, AR coated | ACL12708U-A | 1 | $31.77 | $31.77 | |
| **Light path** | Collimator | Thorlabs | Thorlabs | Fly's Eye Homogenizer, Uncoated, WD=95mm | FLE2 | 1 | $813.96 | $813.96 | |
| | | | | | | **Component Total** | | **$931.63** | |
| **Body** | Base | Thorlabs | Thorlabs | Aluminum Breadboard 24" x 36" x 1/2", 1/4"-20 Taps | MB2436 | 1 | $793.11 | $793.11 | |
| **Body** | Support | Thorlabs | Thorlabs | 95 mm Construction Rail, Clear Anodized, L = 750 mm | XT95-750 | 1 | $220.11 | $220.11 | |
| **Body** | Support | Thorlabs | Thorlabs | Base Plate for 95 mm Rails | XT95P3 | 2 | $58.82 | $117.64 | |
| **Body** | Support | Thorlabs | Thorlabs | 95 mm Rail Construction Clamp, 1/4"-20 Locking Screw | XT95P13 | 1 | $122.28 | $122.28 | |
| **Body** | Support | Thorlabs | Thorlabs | Fixed Arm, Internal SM2 Threads, 60 mm Cage Compatible | CSA1002 | 2 | $351.72 | $703.44 | |
| **Body** | Bracket | Thorlabs | Thorlabs | Right-Angle Bracket for 50 mm Rails | XE50A90 | 1 | $74.27 | $74.27 | |
| **Body** | Tube | Thorlabs | Thorlabs | SM2 Lens Tube, 0.5" Thread Depth | SM2L05 | 1 | $28.42 | $28.42 | |
| **Body** | Light Stop | Thorlabs | Thorlabs | SM3 Ring-Actuated Iris Diaphragm (Ø2.5 - Ø50.0 mm) | SM3D50D | 1 | $156.99 | $156.99 | |
| **Body** | Adapters | Thorlabs | Thorlabs | Adapter with External SM2 Threads and Internal SM3 Threads | SM3A2 | 1 | $35.35 | $35.35 | |
| **Body** | Adapters | Thorlabs | Thorlabs | Male SM2 to Female F-mount Ring | SM2NFMA | 1 | $108.41 | $108.41 | |
| **Body** | Adapters | Thorlabs | Thorlabs | Male M54x0.75 to Female SM2 | SM2A28 | 3 | $34.17 | $102.51 | |
| **Body** | Adapters | Thorlabs | Thorlabs | Male SM2 to Male SM2 | SM2T2 | 2 | $41.29 | $41.29 | |
| **Body** | Fasteners | Thorlabs | Thorlabs | 1/4"-20 Cap Screw and Hardware Kit | HW-KIT2 | 1 | $139.58 | $139.58 | Only a subset required for build |
| | | | | | | **Component Total** | | **$2,643.40** | |
| | | | | | | | **Subtotal** | **$19,155.78** | |
| **Light Source** | Illumination | Lumencor | Lumencor | SPECTRA III Light Engine | 90-10508 | 1 | $22,585.50 | **$22,585.50** | |
| | | | | | | | **Grand Total** | **$41,741.28** | |

**Table S1.** Build list of the transfluorescence macroscope imaging system described herein.
